# Bazam: A rapid method for read extraction and realignment of high throughput sequencing data

**DOI:** 10.1101/433003

**Authors:** Simon P Sadedin, Alicia Oshlack

**Affiliations:** Bioinformatics, Murdoch Children’s Research Institute, Royal Children’s Hospital, Flemington Road, Parkville, Victoria 3052 Australia; Victorian Clinical Genetics Services, Royal Children’s Hospital, Flemington Road, Parkville, Victoria 3052 Australia; Department of BioScience, University of Melbourne, Parkville 3050, Australia

## Abstract

**Background:** As costs of high throughput sequencing have fallen, we are seeing vast quantities of short read genomic data being generated. Often, the data is exchanged and stored as aligned reads, which provides high compression and convenient access for many analyses. However, aligned data becomes outdated as new reference genomes and alignment methods become available. Moreover, some applications cannot utilise pre-aligned reads at all, necessitating conversion back to raw format (FASTQ) before they can be used. In both cases, the process of extraction and realignment is expensive and time consuming.

**Findings:** We describe Bazam, a tool that efficiently extracts the original paired FASTQ from reads stored in aligned form (BAM or CRAM format). Bazam extracts reads in a format that directly allows realignment with popular aligners with high concurrency. Through eliminating steps and increasing the accessible concurrency, Bazam facilitates up to a 90% reduction in the time required for realignment compared to standard methods. Bazam can support selective extraction of read pairs from focused genomic regions, further increasing efficiency for targeted analyses. Bazam is additionally suitable as a base for other applications that require efficient paired read information, such as quality control, structural variant calling and alignment comparison.

**Conclusions:** Bazam offers significant improvements for users needing to realign genomic data.

## Background

The wide scale adoption of high throughput genomic sequencing instruments over the last ten years has generated vast quantities of genomic data with enormous potential for future use. Genomic data is often stored and exchanged as aligned reads in coordinate-sorted BAM or CRAM format. This format is common because many applications (such as viewing the alignment or routine variant calling) can utilise it directly. Storage in aligned form, however, has the significant disadvantage that the data is tied to the reference genome and alignment method used. Many results are highly sensitive to these parameters, and combined data sets typically cannot be analysed together at all unless these parameters are identical. Consequently, to make optimal use of data, users often need to realign the data to a recent genome build and reference. This is resulting in a widespread and growing need for the capability to efficiently realign genomic data.

Realignment of paired reads from aligned data is however both computationally expensive and inconvenient using standard methods. The challenges arise because aligners must access both reads of a pair simultaneously in order to optimally align them. While both reads are usually stored in an alignment file, in a coordinate sorted file a significant fraction may be distant from each other. In these cases, an expensive random lookup is necessary to read the mate information so that both reads of the pair can be written to the output together. Consequently, the standard practice for realignment involves first extracting all the reads, and then sorting them by read name on disk prior to realignment. While this makes extraction feasible, the process is lengthy and requires substantial resources due to the intermediate steps. Interestingly, Picard Tools [1] offers an alternative method to extract read pairs, in the form of SamToFastq, which avoids the need for these intermediate steps in extracting read pairs. However, this method is not widely used in the community. This is likely because SamToFastq is poorly optimised for memory use, making it impractical for use with large data sets. Additionally, Picard Tools cannot target a specific locus, and can only emit a single output stream, causing the process to be bottlenecked by the maximum throughput of a single downstream process (such as alignment).

Here we introduce Bazam, an alternative to SamToFastq that optimises memory use, while offering increased parallelism and other additional features. Bazam increases parallelism by splitting the output streams into multiple paths for separate realignment. Using this technique, a single source alignment can be realigned using an unlimited number of parallel aligners, significantly accelerating the process when a computational cluster or cloud computing resource is available.

**Figure 1.**
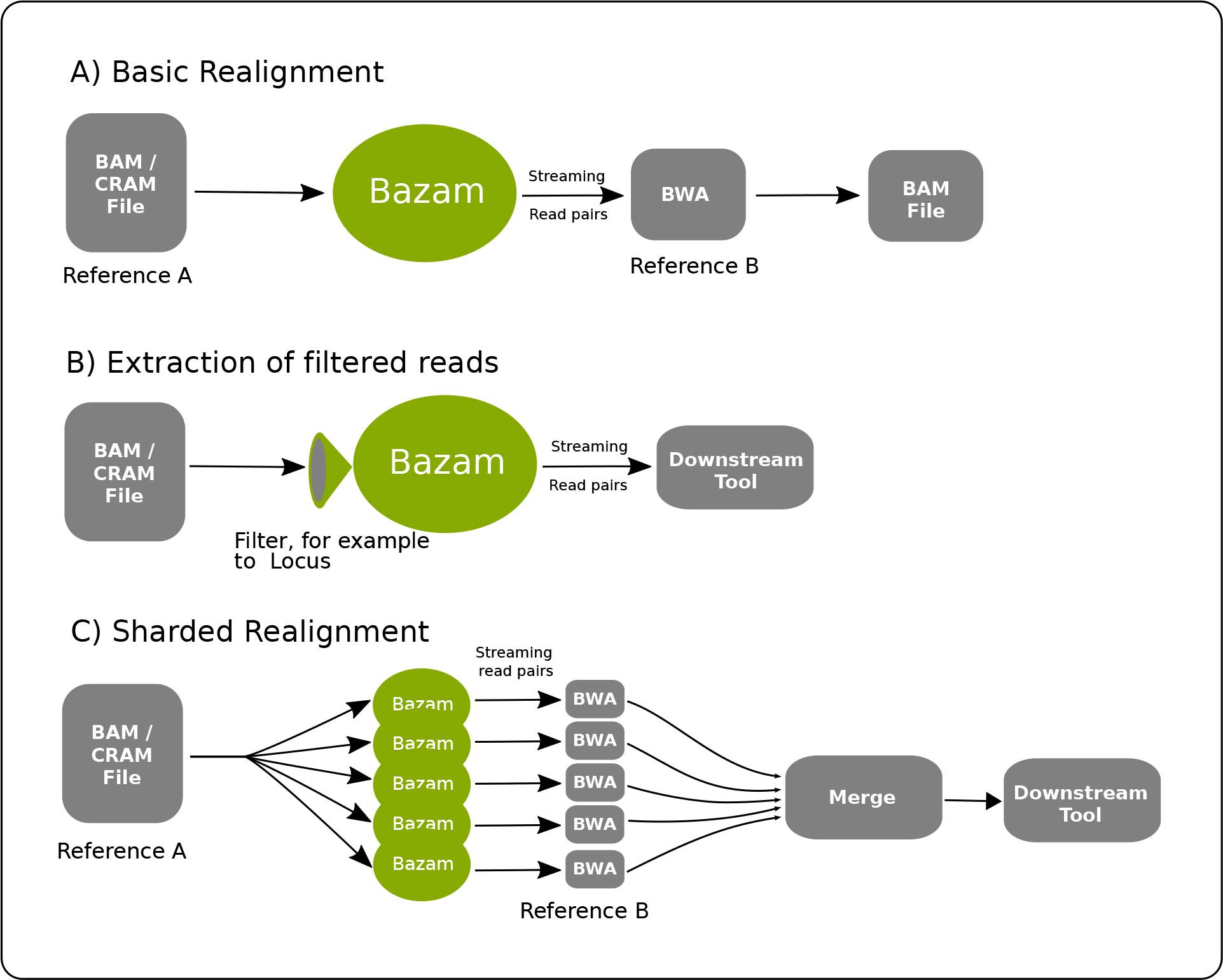
Different configurations for using Bazam. (A) Simple realignment from one reference genome to another without intermediate storage or steps. (B) Extraction of filtered reads such as those overlapping a specific locus. Reads can be streamed to downstream tools directly, or stored in FASTQ format for further processing. (C) Sharded realignment allows for many copies of the aligner to run on different subsets of the data, greatly speeding up realignment.

While realignment is a key application, Bazam also offers utility for any other application relying on detailed read-pair information. Example applications include quality control and structural variant calling. Bazam offers two additional features of particular interest: read position tagging, and localised extraction. Read position tagging renames reads as they are streamed to include the original alignment position in the name of each read. This feature allows ready comparison between new and old alignment positions after realignment. Localised extraction allows realignment to be limited to reads overlapping specified genomic coordinates. Like realignment, this can be achieved using standard tools. However, these tools do not emit both reads of a pair if only one overlaps a region of interest, and are therefore unsuitable for applications that require both of the reads. Here we describe the implementation of Bazam, and demonstrate that it increases efficiency without compromising accuracy.

## Methods

### Pairing of Reads

The primary challenge in extracting paired reads from BAM and CRAM files arises from the predominant choice of coordinate sorted ordering for their storage. This format is used because it places all the reads aligned to a given genomic locus in close physical proximity within the file, maximising efficiency for any analysis focused on short range variation (such as SNV and indel calling, or visualisation in genome browsers). However, coordinate ordering is highly suboptimal for realignment, because a small but significant fraction of reads are located a large genomic distance from their mate. Consequently, a simple linear scan cannot readily extract both a read and its mate in many cases. One possibility is to retrieve each mate as needed using a random seek within the file to the location of its mate. This strategy is highly inefficient, however, because reads are stored within BAM and CRAM files in a block format such that extracting a single read requires decoding some or all of the other reads from the same block. In practice, such a random seek strategy slows down read extraction by several orders of magnitude.

Bazam retains the efficient linear scan of standard methods. However, instead of performing random lookup of each mate, Bazam stores each read in memory until the mate is encountered naturally. For the majority of pairs, both reads derive from the same biological fragment, which is typically closely matched to the reference genome and therefore a short distance on the genome. In these cases, the mate for a read is encountered soon after the read itself, so that the first read needs to be only briefly stored in memory. Reads aligned at a greater distance from their mate must be buffered for significantly longer. Consequently, Bazam requires enough memory to run such that it can store these reads until their mates are encountered by the linear scan. To reduce the memory load, Bazam does not store the full read data structure in memory when the desired output is in FASTQ format. Instead, Bazam stores only data essential to the FASTQ output. Bazam additionally encodes the in-memory reads to compress the data and reduce memory load.

The worst case scenario is represented by a small proportion of reads where the mate aligns to a different reference contig (or chromosome). These reads may represent real structural variation within the sample, but can also be generated artefactually in the preparation of sequencing libraries. In these cases the mate does not resolve at all until its chromosome is encountered by the linear scan. Accordingly, Bazam requires enough memory to store this small proportion of reads for the full duration of the extraction.

By buffering reads, Bazam trades memory for speed. The peak memory required depends on the coverage depth of the alignment, the typical span between paired reads (the insert size distribution) and also on the number of reads whose mates align to different contigs. We observe on typical human whole genome data sequenced at 30x mean coverage depth, that Bazam requires approximately 16 - 32GB of RAM. For cancer genomes or other scenarios with many genomic rearrangements this could potentially increase. However, for many common scenarios the memory requirement of Bazam remains well within the limitations of the resources available in most modern computing systems.

### Parallelism and Sharding

As computational performance is one of the main goals of Bazam, it is designed with a high level of parallelism internally so that system input/output (I/O) is never blocked. This is achieved by using separate threads for reading the input alignment file, writing the output, and a pool of threads to index and buffer each read so that it can be paired with its mate. To further ensure that performance of Bazam can be scaled, read pairs can be split into multiple streams, which is referred to as “sharding”. In sharded mode, several copies of Bazam are run, with each copy emitting a different subset of the reads. Bazam utilises the unique read name assigned to each read pair to ensure that the output streams receive mutually exclusive subsets. Specifically, the name of each read is used to generate a hash code and the modulus of this hash code with the total number of shards is used to decide whether a read is processed by a given Bazam instance. Many copies of Bazam can then run on the same alignment file simultaneously, with each one outputting a unique read subset. This arrangement both reduces the peak memory load and increases parallelism, as each shard can be streamed into different instance of the aligner. With a computational cluster or cloud computing facilities, almost unlimited parallelism is achievable with this method. Sharding can also be utilised to downsample data to lower coverage, by omitting one or more output streams from alignment.

### Software Implementation

Bazam is implemented using Groovy, a modern language derived from Java and which shares most properties with Java including platform independence and very high performance. Bazam uses HTSJDK (https://github.com/samtools/htsjdk) for the underlying BAM and CRAM parsing operations. To enable high concurrency, Bazam employs actor-based concurrency based the GPars framework (http://www.gpars.org/).

## Results

### Efficiency of Realignment

To test the efficiency of Bazam we applied it to a public whole genome data set (NA12878, 30x mean coverage) released as part of the Genome in a Bottle project [2]. First, we realigned the data set from GRCh37 to GRCh38 using both Bazam and using the standard approach without Bazam. The standard approach consists of first sorting reads using `samtools bamshuf`, then extracting them using `samtools bam2fq`, and finally realigning using BWA mem [3], and re-sorting the output BAM file using `samtools sort`. To avoid directly storing intermediate files, this process was constructed using Unix pipes. However, we note that the intermediate sorting stages still write intermediate files, resulting in substantial storage requirements. We refer to this process as Sort-Extract-Realign (SER). In this process we used 16 cores in total as we observed empirically that on our test systems, relatively little improvement in performance was gained by adding additional cores. Picard SamToFastq was run using 32GB of RAM and given 16 processor cores. However in this configuration it failed to complete as it exceeded the allocated memory early in the process. When increased memory was given, anomalies within the data set caused it to abort the process, preventing detailed measurement of its performance. The Bazam process consisted of Bazam directly streaming reads into BWA mem, followed by re-sorting with `samtools sort`.

When using a single instance of BWA, Bazam decreased the overall time by 18.3% and the storage required by 75.9%. In this case, the process was limited primarily by the speed of alignment rather than Bazam’s ability to process reads. When run in sharded mode, however, Bazam was able to split reads between 10 copies of BWA, resulting in a time saving of 91%, while still reducing the storage needed by 63.8%.

**Table 1.** Comparison of run time, memory and storage space between Bazam and a conventional process for realignment

| Tool | Storage Used | Memory | Effective Cores | Time |
| --- | --- | --- | --- | --- |
| Sort-Extract-Realign | 282GB | 20GB | 16 | 13 hours, 15 minutes |
| Picard SamToFastq | - | >32GB | 16 | Did not complete |
| Bazam (no sharding) | 68GB | 29GB | 16 | 10 hours, 41 minutes |
| Bazam 10-way sharding | 102GB | 20GB | 160 | 1 hours, 11 minutes |

### Accuracy of Realignment

We tested the fidelity of Bazam’s read extraction process by comparing Bazam’s output to the expected output using two different methods. First, we converted all reads from the evaluation data set to FASTQ format using the SER method. Then, we aligned these reads to GRCh37 using BWA mem, and re-extracted to FASTQ format using Bazam. Comparison of the two FASTQ data sets found that reads were identical, showing that Bazam reproduces FASTQ with perfect fidelity.

To investigate any unexpected effects resulting from realignment with Bazam, we first realigned the SER-extracted FASTQ to GRCh37, to create an updated alignment using our local alignment configuration. Next, we realigned this updated alignment, with Bazam. These steps ensured that both alignments with and without Bazam used identical reference genomes and aligner settings, so that these factors did not cause artefactual differences.

We then compared the alignments with each other, by applying Bazam’s read position tagging feature. The feature alters read names during realignment to carry the original alignment position. In this way, reads in the new alignment could be readily checked against their old position to identify reads that “moved”.

The comparison between the Bazam and updated realignments revealed a total of 13.7m (1.7%) reads that changed position after Bazam realignment. We hypothesised based on previous studies [4] that this may be caused by ambiguously positioned reads aligning differently due to altered input order. Consistent with this hypothesis, we identified that of the repositioned reads, 92.8% had mapping quality of 30 or less, suggesting their alignments are subject to significant ambiguity. We investigated the moved reads that had high mapping quality and observed that many of the these were mapped to repeat masker regions (Supplementary Table 1), and in many cases were in fact subject to ambiguity despite receiving high mapping quality from BWA. Based on these results, we concluded that Bazam realignment has minimal effect on reads with unambiguous mapping positions, and while reads with ambiguous positions may be repositioned, this is likely due to known behavior of BWA, rather than Bazam itself.

### Application to Repeat Expansion Calling

As an example of Bazam’s utility for aiding downstream analysis tools such as complex variant calling we applied Bazam to STRetch [5], a method for detection of short tandem repeat expansions (STRs) in genomic data. The first step in STRetch selects reads aligning to more than 400,000 known STR repeat regions (as well as any unmapped reads) and then realigns these reads to artificial decoy sequences containing short tandem repeats. When run on pre-aligned data, STRetch extracts all reads within 800bp of each known STR region. This window is chosen to be wide enough to capture both reads of the majority of pairs that fall into the STR region. Nonetheless, some pairs are mapped widely enough apart that they may be missed. We replaced this implementation with Bazam’s local extraction feature and tested the accuracy and efficiency.

When run using the default read extraction method on the same whole genome sample, STRetch took 6 hours and 7 minutes. The unsharded Bazam method reduced the time required to 2 hours and 27 minutes. This improvement is achieved partly by avoiding intermediate FASTQ extraction, but also by eliminating the additional window required for scanning of candidate STR reads. Bazam makes the expanded window unnecessary because it guarantees to output both reads of a pair, even if only one overlaps the extraction window, demonstrating the utility of the localised extraction feature. When run using sharded mode with 6 copies of BWA, STRetch finished in 1 hour and 24 minutes. STRetch primarily derives its sensitivity from its ability to align reads from STR regions to the decoy sequences. Hence we compared STRetch performance between Bazam and standard alignment methods by counting the reads that were aligned to each decoy sequence. We found that Bazam was able to align 3.4% more reads to the decoy sequences than the standard alignment process. Therefore we conclude that alignment using Bazam increases both speed and accuracy in the case of STRetch.

### Wider Applications

While we have primarily developed Bazam with realignment in mind, any application where paired reads are needed can benefit. In particular, we note that many algorithms for complex and structural variant calling are highly dependent on read pair information and hence could benefit from building on this method. Quality control statistics derived from read pair information can also be calculated more efficiently using Bazam than standard methods. Finally, the ability to tag read names with previous alignment information is also useful for benchmarking and comparing alignment software.

## Conclusion

Bazam offers a simple, yet effective tool that enables a significant increase in efficiency and decrease in time required to realign existing genomic data. This has widespread practical utility as the need to reprocess data onto new genome builds with updated alignment software is becoming increasingly prevalent. Bazam also has many other potential uses for applications where full read pair information is needed, especially where extraction from localised regions of the genome is of interest. Bazam is open source software and is available at https://github.com/ssadedin/bazam.

## Acknowledgements

We wish to thanks Cas Simons for helpful discussions about workflows and comments on the manuscript. We wish to thank Harriet Dashnow for comments and explanations regarding STRetch output and software.

